# Prediction Of The Impact Of Genetic Variability On Drug Sensitivity For Clinically Relevant EGFR Mutations

**DOI:** 10.1101/2022.04.25.489389

**Authors:** Aristarc Suriñach, Adam Hospital, Yvonne Westermaier, Luis Jordà, Sergi Orozco-Ruiz, Daniel Beltrán, Francesco Colizzi, Pau Andrio, Robert Soliva, Martí Municoy, Josep Lluís Gelpí, Modesto Orozco

## Abstract

Mutations in the kinase domain of the Epidermal Growth Factor Receptor (EGFR) can be drivers of cancer and also trigger drug resistance in patients under chemotherapy treatment based on kinase inhibitors use. *A priori* knowledge of the impact of EGFR variants on drug sensitivity would help to optimize chemotherapy and to design new drugs effective against resistant variants. To this end, we have explored a variety of *in silico* methods, from sequence-based to ‘state-of-the-art’ atomistic simulations. We did not find any sequence signal that can provide clues on when a drug-related mutation appears and what will be the impact in drug activity. Low-level simulation methods provide limited qualitative information on regions where mutations are likely to produce alterations in drug activity and can predict around 70% of the impact of mutations on drug efficiency. High-level simulations based on non-equilibrium alchemical free energy calculations show predictive power. The integration of these ‘state-of-the-art’ methods in a workflow implementing an interface for parallel distribution of the calculations allows its automatic and high-throughput use, even for researchers with moderate experience in molecular simulations.

## INTRODUCTION

EGFR (Epidermal Growth Factor Receptor) overexpression or mutation can lead to impaired activation of its tyrosine kinase (TK) activity, triggering the hallmarks of cancer, i.e., increased proliferation and survival, aggressive invasion and metastasis, evasion of cell death, and increased metabolism^1,2^. Unregulated EGFR over-activity is present in many types of cancers, especially in those with the poorest prognosis. For example, over 60% of the patients with metastatic non-small-cell lung cancer (NSCLC) overexpress EGFR^3,4^, and around 10-30% of NSCLC patients^5–7^ display mutations leading to ligand-independent activation of the EGFR TK. Some of these activating mutations are located in the extracellular domain and drive a conformational change from the inactive to the active state, mimicking the one induced by the natural ligand (the epidermal growth factor^8^); many others are located at the kinase domain and trigger changes in the ATP-binding pocket altering the ‘on-off’ equilibrium of the enzyme^9–13^ probably by destabilizing the ‘inactive state’ with respect to the ‘active one’^14,15^.

Current treatments for cancers involving EGFR dysregulation are based on either monoclonal antibodies directed against the cognate ligand EGF or EGFR’s homo- or heterodimerization or inhibitors of the tyrosine kinase activity. Various structurally related FDA-approved small molecules either reversibly or irreversibly compete with the natural substrate ATP to inhibit TK activity. Several of these drugs are used in clinics for the treatment of non-small cell lung, pancreatic, colorectal, head and neck, and breast cancers^11,16,17^. The first TK inhibitors (TKIs) with clinical benefits were **Erlotinib** and **Gefitinib**^18–20^, followed by **Erlotinib** (Tarceva®)^21^, **Lapatinib** (Tyverb®^22^), **Gefitinib** (Iressa®)^23^ and third-generation **osimertinib**^24,25^, which forms a covalent bond with C797 after an initial non-covalent binding. Unfortunately, while they show good antineoplastic activity at the beginning of the treatment, as cancer progresses and tumor cells accumulate mutations, drug resistance appears. This is triggered by the emergence of inactivating mutations^26^, which temporarily or permanently rescue the ‘dysregulated’ activity of the EGFR^27–32^. Suggested drug-inactivating mechanisms include, among others: the activation of alternate proteins downstream of EGFR signalling, the activation of proteins that feed into the EGFR signalling cascade, or the decrease in the affinity of the TKIs^33^.

While resistance driven by the rewiring of cellular networks is independent of the molecular details of the EGFR inhibitor and can be tackled by multidrug therapies^34^, resistance due to mutations with decreased inhibitor affinity are dependent on fine details of the drug and the mutation, and are susceptible to theoretical predictions by means of simulation methods. We will focus here on mutations in the kinase domain, which can affect inhibitor binding in a drug-specific manner. In fact, first-generation ATP-competitive inhibitors quickly faced TK mutation-induced treatment resistance related to a decrease in inhibitory potency, which motivates the development of second- and third-generation inhibitors^35^. Unfortunately, while improving resiliency to mutations, second-generation inhibitors display limited efficacy in circumventing some mutations, and even third-generation inhibitors are susceptible to inactivating mutations affecting the vicinities of the ATP binding site^36^.

Improvements in EGFR-TKI based therapies would require detailed knowledge of the impact of mutations on the binding of the drugs. Patient genotyping^37^ followed by *in silico* predictions and *in vitro* validation, could help oncologists ascertain whether mutations render the kinase domain (KD) of EGFR resistant to therapeutic drugs. Furthermore, *in silico* prediction of the impact of mutations in kinase inhibition would not only allow understanding the impact of known mutations on already existing drugs, but would help to anticipate resistance and implement modifications in the therapy before relapse happens. Even more excitingly, *in silico* mutagenesis and binding predictions would allow pharmaceutical laboratories to anticipate inactivating mutations for a drug candidate before it reaches the market, helping to stratify the patient cohorts in clinical trials and triggering the development of a modified drug candidate able to escape from inactivating mutations. To this aim, a reliable simulation-based pipeline has to be developed and implemented in a user-friendly manner for non-experts.

We present here a multilevel and automatized approach that allows the *in silico* prediction of the effect of mutations in the binding properties of TKI targeting EGFR. A variety of sequence-based methods helped to detect some trends of drug-affecting mutations but are far to have any predictive power. Simulation methods explicitly accounting for the structural and dynamic properties of the protein and of the drug-protein complex achieve predictive power ranging from 70% for the lower-level methods to an impressive 100% success for the most elaborate methods based on molecular dynamics and non-equilibrium free energy calculations. We present a workflow based on a recently developed technology that allows a non-expert user to perform complex and sophisticated simulations on the day-time scale with moderate computational resources, hours with a pre-ExaScale parallel supercomputer and minutes in an ExaScale one.

## METHODS

### Dataset

Considerable data on somatic cancer mutations have been gathered via sequencing projects. For this study, sequence variants found in cancer patient samples were extracted from the International Cancer Genome Consortium^38^ (ICGC; https://dcc.icgc.org/), the Catalog Of Somatic Mutations In Cancer^39^ (COSMIC; https://cancer.sanger.ac.uk/cosmic), and the Clinical Variants^40^ (CLINVAR; https://www.ncbi.nlm.nih.gov/clinvar/) databases. For EGFR, the ICGC database yielded 5710 somatic mutations, out of which 5158 are substitutions, including 402 missense substitutions. Only variants leading to changes in the anticancer activity of TKI drugs were retained (Table 1). We searched the literature for the annotated resistance origin (Table S1), and mapped the key regions for binding and activity onto the structure of the TK domain, comprising the nucleotide binding loop (P-loop), the catalytic loop (C-loop), the αC-helix at the dimerization interface, the activation loop (A-loop), the hinge, and the DFG triad (see Figure 1).

**Table 1.** Mutations impacting drug activity and estimates of the effect based on interaction energy profiles (3^rd^ column), PELE docking (4^th^ column) and molecular-dynamics based free energy simulations (5^th^ column); for the latter, the predicted change in free energy of binding is included in kJ/mol. The experimental annotation of the effect of the mutation in drug activity is shown in the last column. In all cases, grey cells indicate prediction errors.

| Mutation | Drug | Eprof (pred) | PELE* (pred) | $\Delta\Delta G(\text{binding})$ FE/MD | Exp. Impact<sup>Δ</sup> |
| --- | --- | --- | --- | --- | --- |
| L718Q | Osimertinib | R | - | 18.7(0.8) R | Resistance<sup>1</sup> |
| G719S | Gefitinib | S | S | -5.5(0.9) S | Sensitive<sup>2</sup> |
| G719S | Icotinib | S | - | -0.9(0.8)<sup>#</sup> S | Sensitive<sup>2</sup> |
| G719S | Erlotinib | S | S | -1.1(0.3) S | Sensitive<sup>3</sup> |
| G719S | Lapatinib | S | S | -5.1(2.8) S | Sensitive<sup>4</sup> |
| L747S | Gefitinib | S | S | 13.8(1.8) R | Resistance<sup>5</sup> |
| L747F | Osimertinib | S | - | 4.4(1.2) R | Resistance<sup>6</sup> |
| L747H | Osimertinib | S | - | 6.5(3.6) R | Resistance<sup>6</sup> |
| S768I | Gefitinib | S | R | -1.8(0.4) S | Sensitive<sup>7</sup> |
| V769M | Gefitinib | S | S | -7.7(2.2) S | Sensitive<sup>8</sup> |
| T790M | Gefitinib | S | S | 15.5(0.9) R | Resistance<sup>9</sup> |
| T790M | Erlotinib | S | R | 17.6(1.9) R | Resistance<sup>9</sup> |
| T790M | Lapatinib | S | R | 19.3(0.9) R | Resistance<sup>10</sup> |
| T790M | Osimertinib | S | - | -0.5(2.1) S | Sensitive<sup>11</sup> |
| T790M | Icotinib | S | - | 11.5(0.6) R | Resistance<sup>12</sup> |
| L792F | Osimertinib | S | - | 4.2(1.0) R | Resistance<sup>13</sup> |
| L792H | Osimertinib | R | - | 8.8(0.1) R | Resistance<sup>13</sup> |
| G796S | Osimertinib | S | - | 4.8(0.4) R | Resistance<sup>14</sup> |
| C797G<sup>&</sup> | Osimertinib | R | R | Resistance | Resistance<sup>15</sup> |
| C797S<sup>&</sup> | Osimertinib | R | R | Resistance | Resistance<sup>16</sup> |
| L833V | Gefitinib | S | S | -5.7(1.4) S | Sensitive<sup>17</sup> |
| H835L | Gefitinib | S | S | -7.5(1.3)<sup>#</sup> S | Sensitive<sup>17</sup> |
| L838V | Gefitinib | S | S | -6.3(3.1) S | Sensitive<sup>18</sup> |
| T854A | Gefitinib | R | S | 13.2(1.1) R | Resistance<sup>5</sup> |
| L861Q | Gefitinib | S | S | -5.4(0.7)<sup>#</sup> S | Sensitive<sup>19</sup> |
| T790M/C797S | Erlotinib | R | R | 6.2(2.4) R | Resistance<sup>20</sup> |

\* PELE calculations require X-ray structures as an input and cannot be used to explore covalent binders.
& A trivial prediction of resistance for covalent inhibitors binding to C797

**Figure 1:**
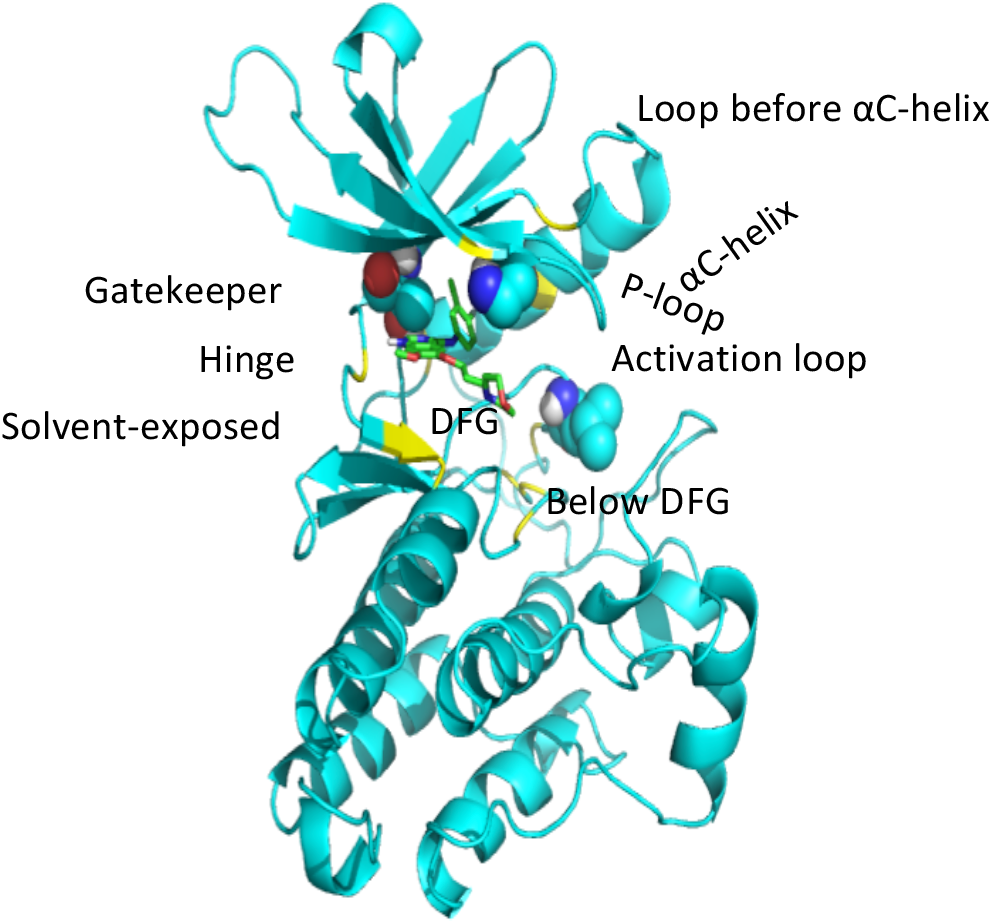
Location of clinically relevant mutations in the kinase domain of EGFR: The secondary structure of EGFR is displayed in cyan, the clinically most mutated residues as balls, and the other mutations, which we consider in yellow. The inhibitor gefitinib is shown as sticks with green carbon atoms (see Table 1).

### Sequence analysis

We used sequence alignments to determine if mutations affecting drug-binding are located in variable or conserved regions and whether or not such mutations are already common even in the absence of the evolutionary pressure of the drug. To this end, we extracted sequences of 94 human tyrosine kinases from KinBase^41^ (http://kinase.com/web/current/kinbase/), aligning them with ClustalW as implemented in the msa R package^42^. Sequence variability at each position of the TK domain was determined from the Shannon entropy score as described elsewhere^43^ (see eq. 1):

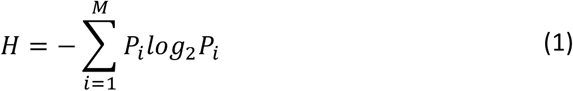

where the sum extends for all mutations sampled in the alignments at a given position, and *P*_*i*_ stands for the probability of the existence of residue i at a given position. Non-pathological human polymorphisms mapping onto TK domain of EGFR were extracted from the gnomAD database^44^.

### Pathogenicity analysis

We used the PMUT program^45,46^ to evaluate the pathological potential of mutations affecting drug response, ranging from sequence and structural information to the protein^47,48^.

### Modelling of the complexes

We use the structure of PDB entry 4WKQ as a template, using PDB entry 2ITY to fill the structural gaps. Both structures contain **Gefitinib** (Iressa®) and cover a large portion of the protein at high resolution (1.85 Å, covering residues 694 to 1020, and a resolution of 3.42 Å, covering residues 697 to 1019). Refinement involved checking alternative conformations per residue in the 4WKQ parts, keeping only the ones with the higher occupancy. For the crucial D855 in the DFG-motif, we chose as a starting point the side-chain orientation making a double salt bridge with the catalytic K745 and E762 in the αC-helix. The final model was subjected to structural checking using our local validation module^49^. The **Erlotinib** binding geometry was taken from 4HJO, the one for **Lapatinib** from 1XKK, and the one for **Ositmertinib** from 4ZAU. The **Icotinib** binding geometry was taken from the *biobb* AutoDock-Vina^50^ module defining the binding pocket as those residues closest to 6.5 Å from **Gefitinib** in the 4WKQ structure. In all cases, structural waters present in the crystal around the binding site were maintained, the hydrogens being oriented during the molecular dynamics (MD) setup procedure as discussed below.

### Force-Field parameters

Proteins were described by the AMBER99SB-ILDN force-field^51^, water by the TIP3P model^52^, and counterions by the AMBER99SB-ILDN associated ion model. For ligands, we used our web-based automatic tool taking care of defining the suitable charge state of the ligands (suitable charge state and parameters were obtained using an automatized protocol^53–55^).

### Generation of mutant starting geometries

Starting from the wild type (WT) geometries above, the *biobb_structure_checking* module was used to create and validate the geometry of the mutants. The structure-checking steps involved: proper amide assignments, chirality, cis/trans backbone, disulphide bridges, and severe intra-protein clashes. Neither the WT nor any of the modelled mutants presented any major issue. In any case, models were subjected to minimization, thermalization, and 50 ns equilibration before production. For each mutant, the residue protonation state was defined at a physiological pH of 7.4 using the *PROPKA* software (v3.1.8)^56^, reorienting side chains of histidines with the PDB2PQR (v2.1.1) webserver^57^ (https://server.poissonboltzmann.org/pdb2pqr).

### Induced fit calculations

We used the PELE suite of programs to analyse the docking of ligands to wild type and mutant proteins in an unbiased manner. We compared the docking energies of the wild type and mutant to detect mutation-induced changes in binding. PELE uses a Metropolis-Monte Carlo/annealing protocol that combines ligand random moves, main chain perturbation based on normal displacements, and a final relaxation stage consisting of side-chain relaxation and global minimization. Crystal structures (see above) for the wild type and mutants were prepared with Schrodinger’s Protein Preparation Wizard^58^ and Maestro^59^, defining a docking sphere of a radius of 5 Å around the ATP binding site. Defaults were used for PELE^60^. OPLS2005^61^ was used to define the energy functional combined with the SGB solvent model corrected for nonpolar contacts^62^. Rotamer libraries were taken from our Peleffy library. Binding poses were obtained using AdaptivePELE^63^ to explore diverse binding poses within the docking sphere, clustering the poses and re-exploring the most relevant ones.

### Molecular Dynamics (MD) simulations

For each apo and holo EGFR variants, ten replicas were generated. Each of them was optimized to relax bad contacts, solvated in dodecahedral water boxes extending for at least 12 Å from any atom of the protein. Counterions (Na^+^ and Cl^-^) were added to maintain neutrality adding additional ions to adjust to a 150 mM salt concentration. Solvated systems were then reoptimized, slowly thermalized (310 K) and equilibrated, first in the NVT ensemble for 1 ns before moving to the NPT one, slowly removing restraints on the protein and the ligand heavy atoms for 1 ns. Each replica of the final systems was relaxed for 50 ns in the NPT ensemble (P=1 atm; T=310 K) before production runs (100 ns each replica), from which seeds for forward and reverse free energy calculations were extracted (see below). Integration of Newton’s equations of motion was done every 2 fs using LINCS^64^ to maintain all bonds involving hydrogen frozen at equilibrium distances. Periodic boundary conditions and Particle Mesh Ewald methods^65,66^ were implemented to capture long-range interactions. Parrinello-Rahman thermostats/barostats^67,68^ were used to maintain pressure and temperature at desired values. All MD calculations were done using GROMACS (v2018.4)^69^.

### Interaction profiles

We used the collected trajectories of the complexes to determine the residue-drug interaction energies and determine those residues with the strongest interactions^70^ as determined from the combination of electrostatic (determined by Poisson-Boltzman calculations^71^) and van der Waals interactions. Group fragmentation was done to maintain monopoles as describe elsewhere^71^.

### Free energy calculations

The mutation-induced change in the binding free energy for the different inhibitors was computed using standard thermodynamic cycles comparing the free energy change associated with the mutation in the apo and drug-bound protein states (Figure 2). Individual free energies were computed via non-equilibrium methods: the *Crooks Gaussian Intersection (CGI), Jarzinsky’s equality (JE)* and the *Bennett Acceptance Ratio (BAR)* methods^72^. Contrary to free energy perturbation or thermodynamic integration (TI), non-equilibrium methods determine the free energy of an alchemical process from the distributions of irreversible work due to a system change (an amino acid mutation in our case), obtained in ‘forward’ and ‘reverse’ directions (see Figure 2). We generated a meta-trajectory concatenating the 10 collected replicas for the WT and mutant in both apo and holo states (see Figure 2), selecting 100 random configurations as starting points for slow-growth TI non-equilibrium perturbations using de Groot’s PMX protocol^73,74^. For each mutation, we run 100×2×2 alchemical changes. Each of the perturbation TI trajectories was extended for 50 ps after a series of test analyses had demonstrated it to be a good compromise between accuracy and computational efficiency (see Suppl. Figure S1). The histograms of irreversible work for the forward and reverse transitions were calculated by the three methods outlined before (see Figure 2) to determine three independent estimates of the reversible free energy associated with the mutation. Where a discrepancy between the three estimates was large (standard deviation of more than 3 kJ/mol, with one of the estimates leading to a global change in the predicted free energy change), simulations were extended to 500×2×2 alchemical changes to check for convergence between the different estimates.

**Figure 2:**
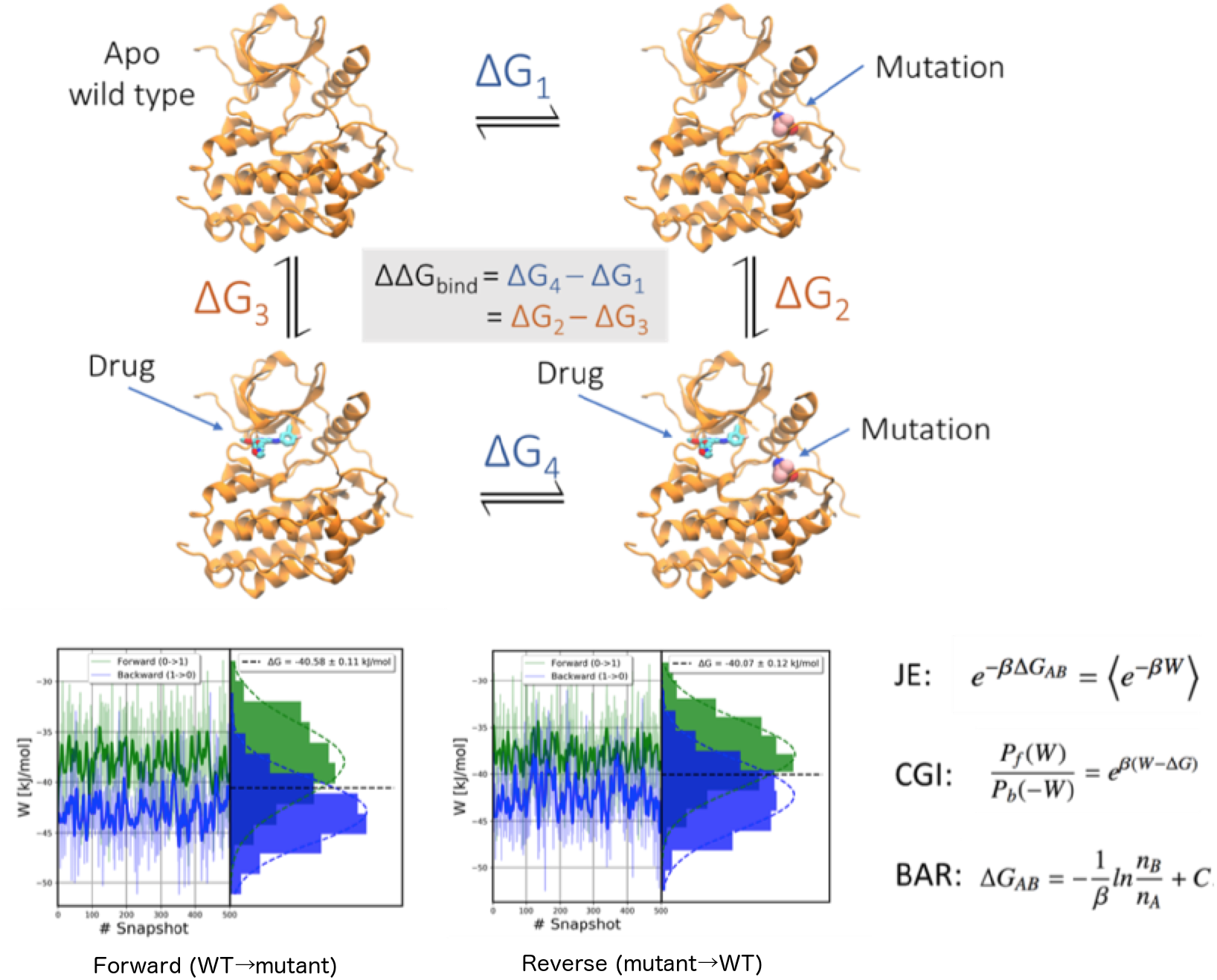
TOP: Thermodynamic cycle used to determine changes in binding free energy associated with protein mutations. BOTTOM: Examples of histograms of works obtained by mutating one residue into another in apo (left) and holo (right) EGFR, respectively. Equations on the right correspond to the three methods considered here to derive free energies for the histograms of irreversible work (Jarzinski equality (JE; top), Crooks Gaussian Intersection (CGI; middle) and Bennett Acceptance Ratio (BAR; bottom). W stands for the reversible work associated to the A → B mutation, P refers to the histograms (forward and reverse), and C is a constant defined from the A and B partition functions (see Methods).

### Free energy workflows

Calculations above imply the use of a myriad of tools. To use them efficiently, they were organized in execution pipelines (workflows) built using the BioExcel Building Blocks library^49^ (abbreviated from here onwards as *biobb*; https://github.com/bioexcel/biobb; https://mmb.irbbarcelona.org/biobb/). A new HPC-focused workflow was specifically developed for this project, taking care of all the mutations, MD setup and PMX calculations, circumventing human intervention to prepare many thousands of individual simulations (see Figure 3). To ensure efficient usage of pre-exascale computational resources, the workflow was launched and controlled using the PyCOMPSs workflow manager^75^, which automatically distributed the workflow individual tasks in a parallel manner in HPC supercomputers. A typical run implies the use of c.a. 768 to 3,072 cores of the MareNostrum supercomputer at the Barcelona Supercomputing Center. The protocol has been tested in PRACE supercomputers showing excellent parallelism on more than 40,000 cores.

**Figure 3:**
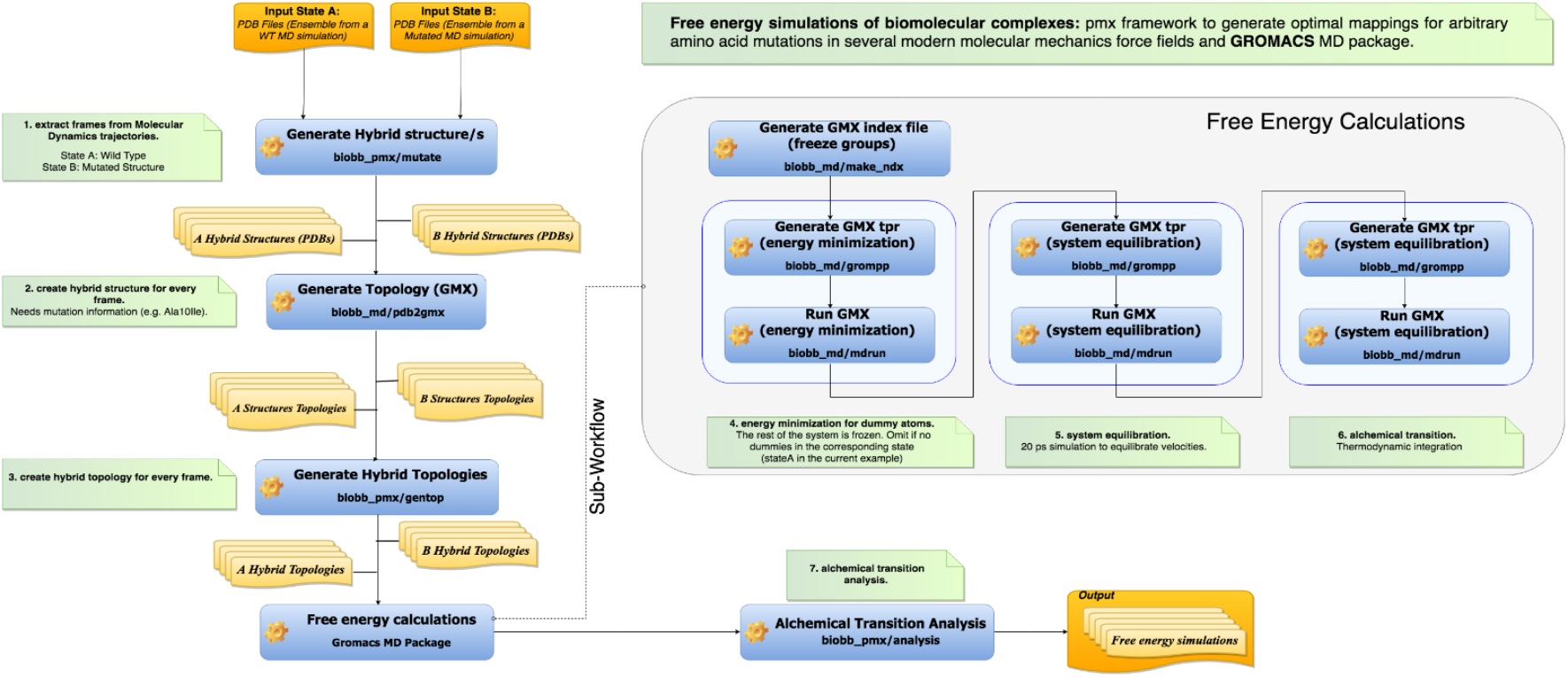
Relative free energy calculation workflow using BioExcel Building Blocks, wrapping GROMACS and the PMX software.

## RESULTS

### Sequence analysis

The ClustalW multiple sequence alignment (Figure 4A) suggests that drug-related mutations tend to be placed at conserved regions (9 out of 14 mutations sites map on regions with Shannon’s score H<2), in some cases in ultra-conserved positions such as or 796 or 835. However, there are drug-affecting mutations mapping on variable positions and there are many highly conserved positions for which no drug-affecting mutations are described. Based on sequence conservation, it is difficult to distinguish between positions were resistance-mutations are detected and those where a mutation induces an equal or better response of the drug. Therefore, multiple alignment analyses do not have enough predictive power to determine which positions are more likely to concentrate mutations affecting drug activity. The analysis of BLOSUM62 matrices^76^ suggests that in general, drug-affecting mutations imply moderate changes in the nature of the amino acid and no disruptive mutation is detected in the list of drug-affecting mutations (Figure 4B). Interestingly, there is no relationship between the drastic change induced by a mutation and its impact on drug-resistance. For example, the ‘disruptive’ change G719S (see Figure 4B) does not lead to resistance, and the ‘mild’ T790M one inactivates most of the TK inhibitors. The inspection of BLOSUM62 matrices does not allow us to predict drug-affecting mutations. Except one, all drug-affecting residue changes can be explained by single nucleotide changes, which suggest that they appear spontaneously as human polymorphisms in the absence of drug pressure. However, this is not the case, as with a very few exceptions, drug-affecting mutations do not appear as human polymorphisms (Figure 4C). Thus, we can conclude that drug-affecting mutations are a consequence of stressed replication in cancer that helps to accumulate mutations. Some of these mutations, but not all, will exhibit positive selection as they lead to better survival of the cancer cells by inactivating drug response.

**Figure 4:**
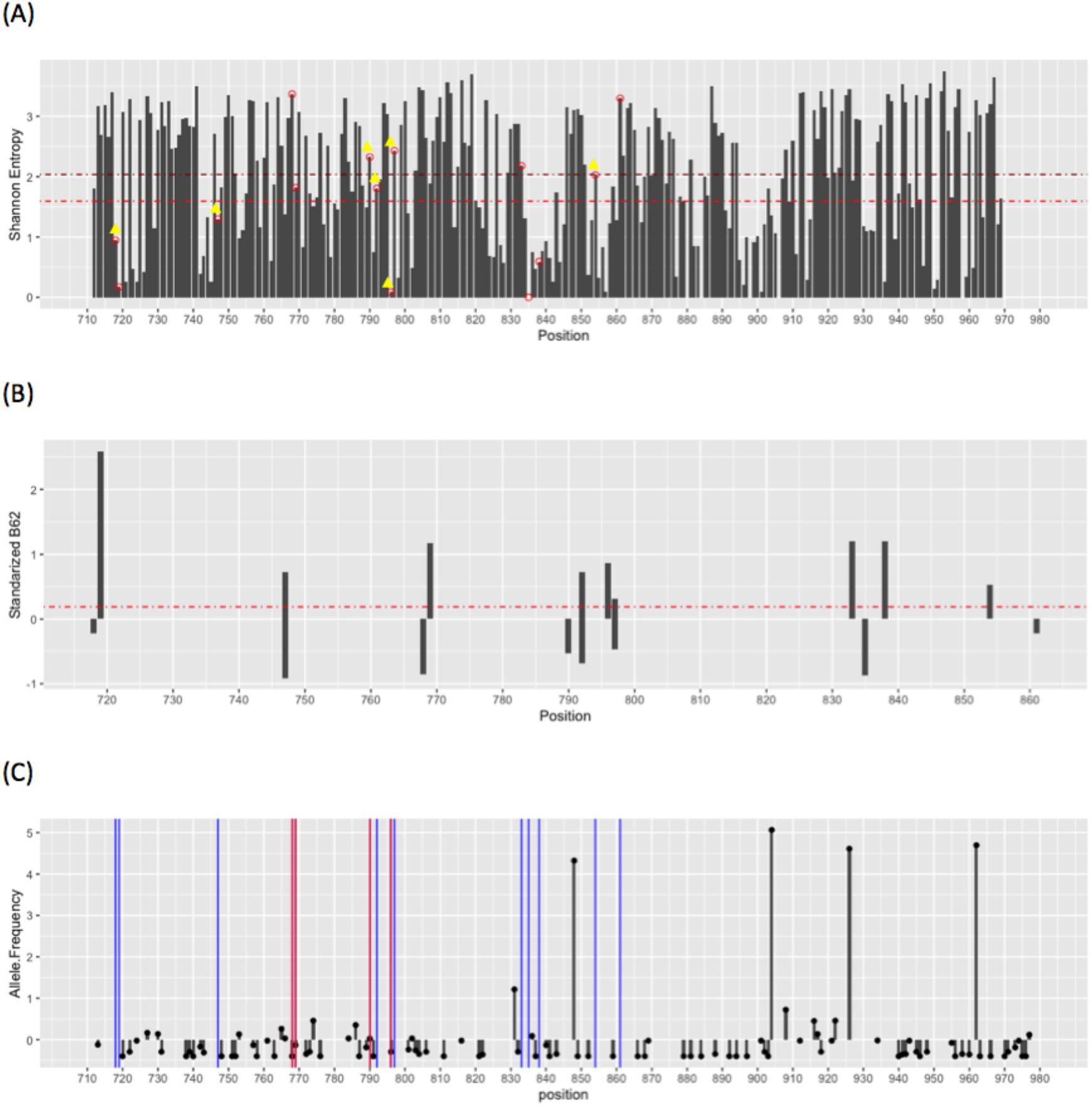
(A) Shannon’s entropy on the human kinase domains (0 fully conserved, 4 fully variable). Places of drug-affecting mutations are marked with circles and those leading to drug resistance are marked with yellow triangles; the horizontal lines correspond to average Shannon’s entropy (see Methods) in the tyrosine kinase subset of the human kinome for all positions (magenta) and those where drug-affecting mutations are found (red). (B) Standardized BLOSUM62 index (referred to the expected ones for random mutation of the wild type residue at this position) associated to the drug-driven mutations, negative values implies that drug-affecting mutations are more aggressive than expected, and positive values the opposite. (C) Standardized allelic frequencies of polymorphisms in the tyrosine kinase domain of EGFR found in the GNOMAD database of human polymorphism. Vertical lines refer to the position of drug-affecting mutations (red when mutation is found as a natural polymorphism; blue when drug-affecting mutation does not map a known human polymorphism).

To complete the sequence analysis, we used PMUT (see Methods) to determine the general pathological potential of positions concentrating on drug-affecting mutations, as well as the specific pathological nature of each drug-affecting mutation. Figure 5A shows that, in general, the TK domain (712-979) is the part of EGFR where a higher profile of pathogenicity is expected from any kind of mutations, which contrasts with the much permissive N- and C-terminal domains. In general, positions concentrating drug-affecting mutations are signalled as ‘pathological positions’ (Figure 5B), but there is not a dramatic difference between the average pathogenicity score of drug-affecting positions and the rest of the TK domain (Figure 5B). Finally (Figure 5C), with one exception (the gatekeeper mutation T790M, corresponding to a polymorphism), the rest of the drug-associated mutations imply a pathogenic risk similar to that of a random mutation mapping the same region. This stands for both mutations leading to an equal or better response of the drug and those inactivating them. In summary, pathological predictions give almost no clue on whether a mutation should have any impact on the activity of the TK inhibitors.

**Figure 5:**
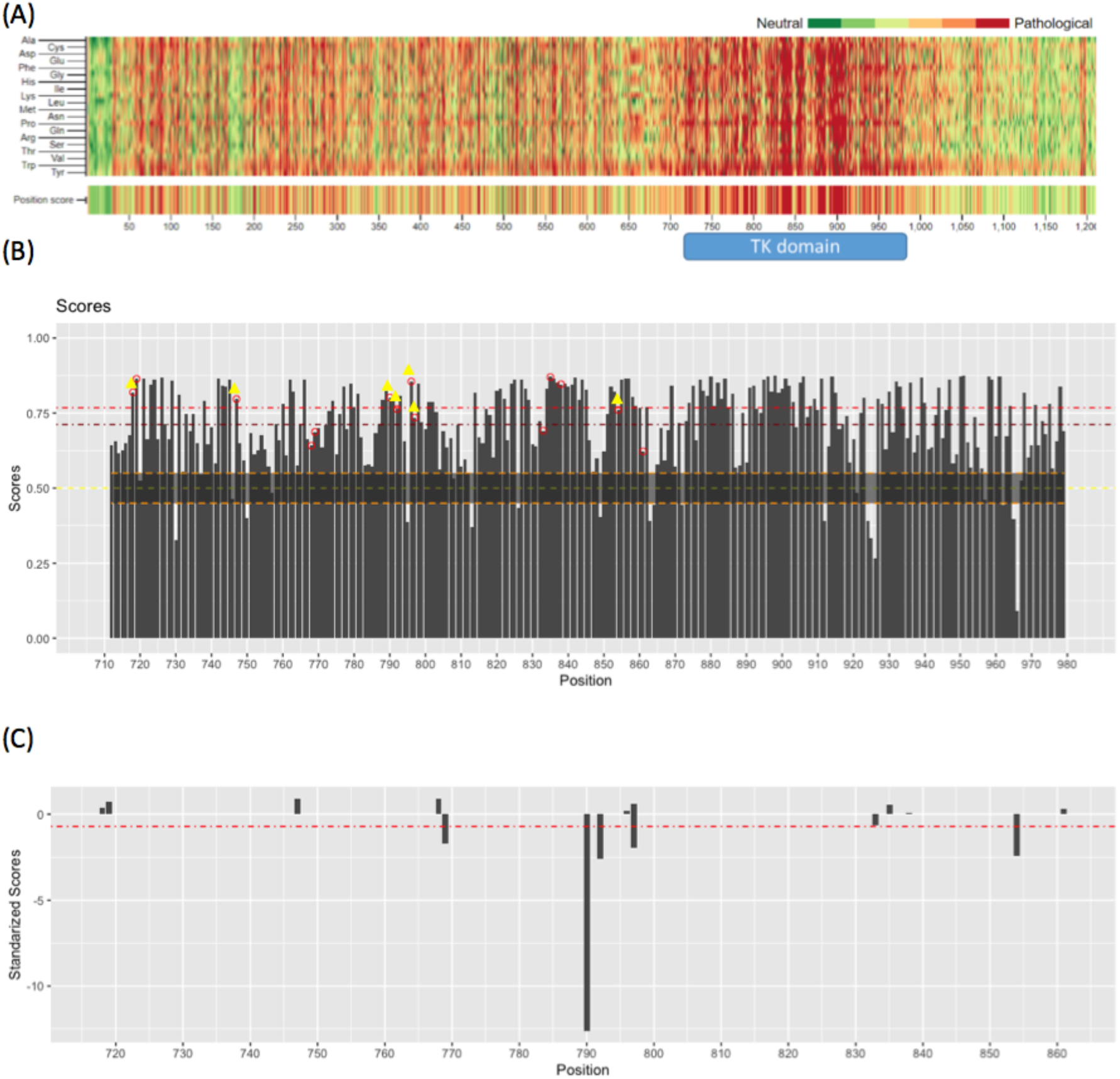
(A) Pathogenicity map of EGFR according to PMUT calculations; the upper part of the plot considers the 19 unique mutations at each position and the bottom part of the plot the average pathological index at this position (the region of the tyrosine kinase domain is highlighted). (B) Average pathogenic profile of the tyrosine kinase domain of EGFR (averaging over the 19 potential mutations). The positions of drug-affecting mutations are marked with small circles, and yellow triangles in the case of mutations generating drug-resistance; the yellow dashed line indicates the criteria to classify pathological (above) or neutral (below) mutations; the magenta dashed line indicates the average pathogenicity index of the tyrosine kinase domain, and the red one that of the positions where drug affecting mutations are detected. (C) Standardized pathogenicity index of the drug-affecting mutations, positive values indicating more pathological than expected and negative values more neutral effect than expected.

### Interaction energy profiles

As sequence-based techniques fail to have predictive power on the impact of mutations on the modulation of the therapeutic response of drugs, we explore the drug-protein interaction profiles (see Methods) obtained from the ensembles collected from MD simulations. Results in Figure 6 provide interesting information on which residues are dominant in ligand-protein interactions. Thus, for the natural substrate ATP, residues involved in ATP or Mg^2+^ binding are the most prevalent for modulating binding (Figure 6 top left panel). They are: i) highly conserved in multiple TK alignments, ii) rarely involved in polymorphisms, and iii) never mutated due to drug treatment; clearly, the pressure to have a functional protein protects these crucial residues from mutations. The interaction profiles between protein and inhibitors are quite different to that of the ATP-complex, but the similarity among them is quite surprising (Figure 6), indicating that all the inhibitors, including the 3^rd^ generation ones, are exploring a similar region of the protein (note that here we explore the previous non-covalent binding before bond formation). There are, however, quantitative differences between the different drugs, especially in terms of the intensity of the interactions with diverse residues. For example, T790 is consistently a stabilizing residue for all the inhibitor binding, except Osimertinib, where its effect is negligible, or L718 whose interaction with Osimertinib is much more favourable than with 1^st^ or 2^nd^ generation inhibitors. Globally (Figure 6), around 80% of drug-resistance mutations happen at positions where wild type residues show favourable drug interactions, while also around 80% of the mutations leading to drug susceptibility are placed at positions where the wild type has neutral or unfavourable drug interactions. However, drug-affecting mutations are rarely located at residues showing very strong interactions with the drug (labelled in black in Figure 6). Thus, interaction profiles provide information on the regions that are prone to concentrate drug-affecting mutations but are unable to predict precisely the position that can mutate.

**Figure 6:**
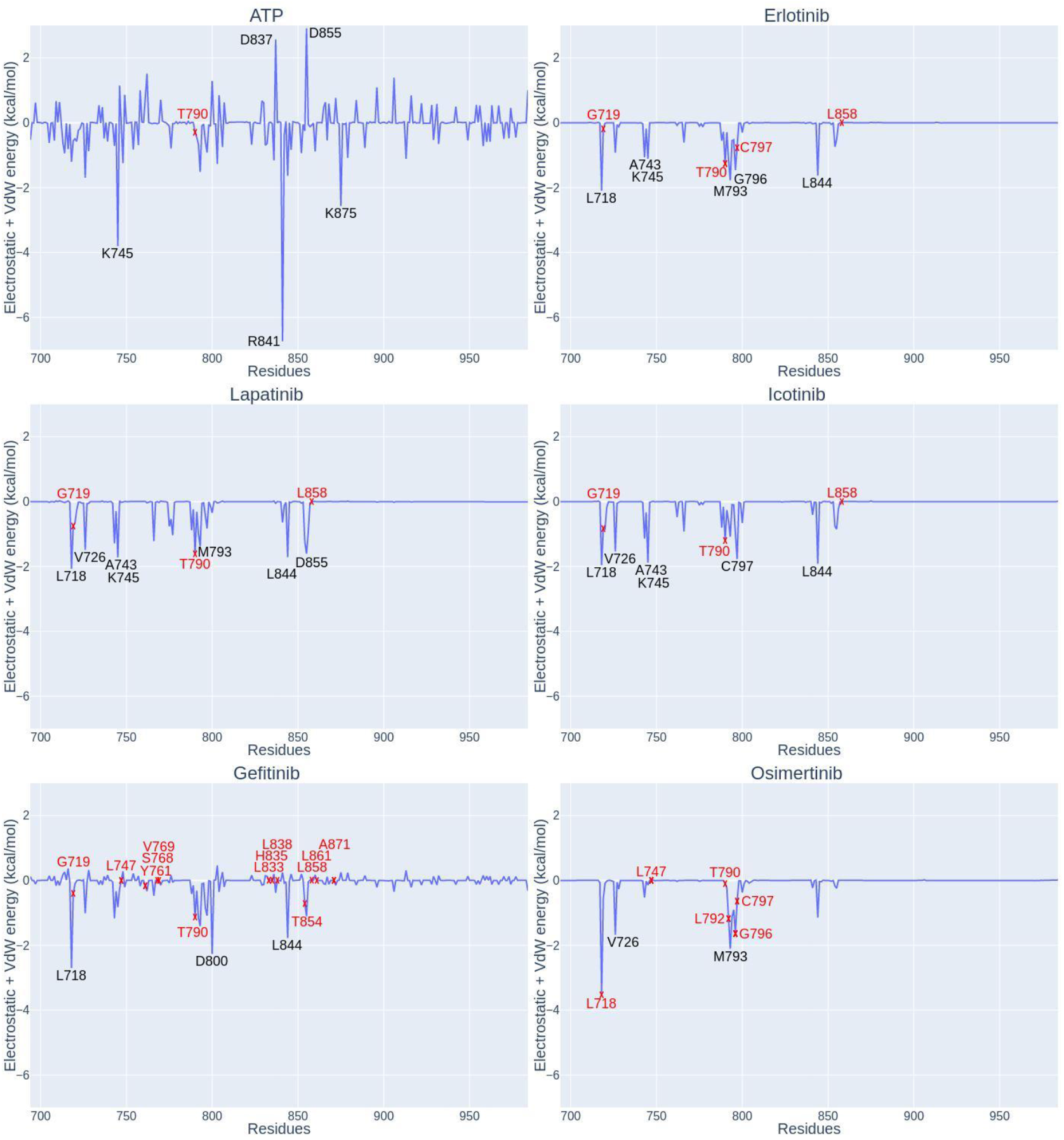
Interaction profiles for natural substrate ATP and inhibitors of the tyrosine kinase domain of EGFR. Residues involved in the interaction with the binder are shown in black and mutations which affect binding are shown in red.

To determine whether the interaction profiles can or cannot predict the impact of the specific mutation (at a given position) on drug activity, we compare MD-derived interaction profiles from wild type and mutant proteins. In general, differential interaction profiles indicate that the protein accommodates the drug-induced mutations well, and the total of drug-protein interactions is not much altered (see Table 1 and examples in Suppl. Figure S2). This result agrees qualitatively with results from the BLOSUM analysis above, confirming that in general drug-affecting mutations are mild. Unfortunately, differential interaction profiles fail to detect some well-characterized resistance mutations, for example, those linked to mutations at L747 and T790. In the first case, interaction profiles do not recognize L747 as a crucial position for stabilizing the drug (Figure 6), and accordingly, mutations to different residues (Ser, Phe or His) are predicted to be innocuous for binding (see example in Suppl. Figure S2). The case of the T790 gatekeeper is different, here the energy profile detects threonine as stabilizing (Figure 6), but the substitution to methionine is not predicted to make such interaction weaker. Therefore, some mutations reducing drug activity are not changing dramatically the direct inhibitor-protein interactions, and the destabilizing effect should be related to structural distortion, solvent bridges or other effects, which are not captured by these simple calculations.

### Exploration of docking landscape

Results above suggest that the impact of some mutations on the binding of inhibitors to the TK domain of EGFR cannot directly be explained by changes in direct interactions between the drug and the protein, and in fact, in some cases they appear at positions that are clearly involved in inhibitor recognition. We explore then docking landscape using PELE (see Methods), which should allow us to detect changes in binding related not only to direct binding, but also to the easiness of drug entrance, or the cost of reorganizing the protein residues at the binding site, aspects that are not considered in simple energy interaction plots. Results in Table 1 show that PELE calculations succeed in predicting drug-induced resistance in 76% of the cases (13 out of 17) compared to 65% of differential interaction energy profiles (17 out of 26). PELE predictions are in general incorrect when water-mediated interactions between the protein and the drug are neglected (Suppl. Figure S3). These interactions are not explicitly considered in PELE calculations and stabilize the interactions of inhibitors with T790 and T854.

### Molecular dynamics-based free energy calculations

These calculations combine equilibrium MD simulations with non-equilibrium alchemical mutations in the *apo* and *holo* states of the protein and represent the ‘state-of-the-art’ in atomistic simulations. Traditionally, its practical use requires thorough expertise and considerable computational resources, but the workflow developed here (Figure 3) allows automation, full use of massively parallel computer architectures and a simple use even for non-experts. The only source of uncertainty arises from the small divergence in the estimates among different integration methods. The analysis of histograms (Figure 2) reveals that problematic cases are typically related to poor overlap between the ‘forward’ and the ‘reverse’ histograms, which generate noise that is maximized in CGI where a Gaussian distribution of irreversible work is assumed. Very encouragingly (Table 1), these divergences are detected in only a few cases and are corrected by simply extending simulations. By construction, FE/MD methods contain all the enthalpic and entropic contributions to binding and a good force-field, combined with appropriate simulation length, should provide accurate estimates. Indeed, the predictive power of the FE/MD protocol outlined here is perfect, as it succeeds in correctly classifying drug-affecting mutations in all studied cases, even in those where simpler methods based on the analysis of differential energy profiles or Monte Carlo PELE calculations fail. We cannot expect this performance to translate to all proteins, drugs, and mutations, but the excellent results obtained by Chodera’s lab on a related system using also alchemical free energy simulations^77^ rises optimism that these sophisticated simulation might be of general use to predict the impact of single point mutations on drug activity, even at preclinical or early clinical stages. The intrinsic complexity of these calculations that limits their use to a bunch of highly expert groups is reduced by the development of robust workflows, whose use do not require expertise and that allows a perfect parallelism allowing to obtain results in a time scale compatible with pre-clinical and clinical use.

## DISCUSSION

Sequence analyses provide useful information on the origin and placement of drug-affecting mutations. In most cases, they are generated by single nucleotide changes and are typically located in conserved regions, where the pathogenic risk associated with mutations is high. Specific positions where drug-affecting mutations happen show a conservation level similar to that of the neighbouring regions. The mutations leading to alteration in drug efficacy tend to be mild in terms of changes in amino acid properties and are not more ‘pathogenic’ than the average expected value at that position. With a few exceptions, drug-associated mutations do not match polymorphisms, which suggests that high stress in replication and most likely poor proofreading of the nascent DNA is required for the appearance of these mutations. Finally, a significant number of the studied mutations imply equal or higher sensitivity to the drug, which means that not all drug-affecting mutations correspond to a canonical positive selection paradigm. Overall, sequence-dependent trends are useful to define regions where mutations are susceptible to altering the response to the drug, but are not able to predict in advance when a mutation will cause resistance to chemotherapy.

Energy profiles detect well which regions establish strong interactions with the ligand and, accordingly, are more informative than sequence analysis to detect precisely the ‘susceptible’ regions where mutations might impact the activity of the drug. However, the success rate is just moderate as there is a non-negligible number of cases where the impact of mutations in ligand binding is modulated by non-direct interacting terms. When flexibility and diffusion considerations are incorporated in the evaluation of drug binding, the predictive power increases, but not dramatically (up to 76%), with cases where we cannot reproduce experimental findings, in most cases due to the involvement of water-mediated interactions that are not easily captured by a method based on continuum solvation models.

Non-equilibrium alchemical free energy calculations provide results of an astonishing accuracy (100% success rate), based only on physical principles without any *ad hoc* training process. By construction, assuming a good force-field and extended sampling, the protocol should capture the different contributions (enthalpic and entropic) to differential binding and has the advantage to provide a physical rationale for the effect of the studied mutations. The limitations of these types of calculations are clear: i) they require user expertise, ii) set-up of the calculations is difficult as it implies thousands of individual simulations each of them requiring several preparation steps, iii) these calculations are computationally expensive and might require very large wall clock times, hampering its practical use in clinical environments. The BioExcel Building Blocks-based workflow developed here allows us to simplify dramatically the complexity of launching simulations, which does not require any specific training in advanced simulation methods. Furthermore, using a clever workflow manager (PyCOMPSs; see Methods) allows extremely fast and efficient parallelism which reduces the wall-clock associated with the entire process to hours when using a pre-exascale supercomputer and most likely minutes in an ExaScale machine. We speculate that once fully calibrated and tested, protocols like the one shown here could be used to accurately predict mutations affecting drug activity already in the *in silico* stages of drug design, helping to develop alternative drugs anticipating inactivating mutations before they appear.

## ACKNOWLEDGEMENTS

We are indebted to BioExcel partners, especially Prof. de Groot’s group for helpful discussion on PMX calculations and Prof. Rosa M^a^ Badia for help with the PyCOMPSs programming model. This work has been supported by the BioExcel-2 Centre of Excellence for Computational Biomolecular Research (823830), the Spanish Ministry of Science (RTI2018-096704-B-100, PID2020-116620GB-I00) and the Instituto de Salud Carlos III–Instituto Nacional de Bioinformática (ISCIII PT 17/0009/0007 co-funded by the Fondo Europeo de Desarrollo Regional). Funding was also provided by the MINECO Severo Ochoa Award of Excellence from the Government of Spain (awarded to IRB Barcelona). M.O. is an ICREA (Institució Catalana de Recerca i Estudis Avancats) academia researcher. The IRB is supported by MINECO Severo Ochoa Award of Excellence from the Government of Spain. Nostrum Biodiscovery is supported by Fundación Marcelino Botí n (Mind the Gap), CDTI (Neotec grant -EXP 00094141/SNEO-20161127) and Torres Quevedo grant (PTQ2018-009992).

## Notes

### Competing Interest Statement

The authors have declared no competing interest.

