## Supplementary Material for "Prediction Of The Impact Of Genetic Variability On Drug Sensitivity For Clinically Relevant EGFR Mutations"

*References to Table 1*

*Supplementary Figures*

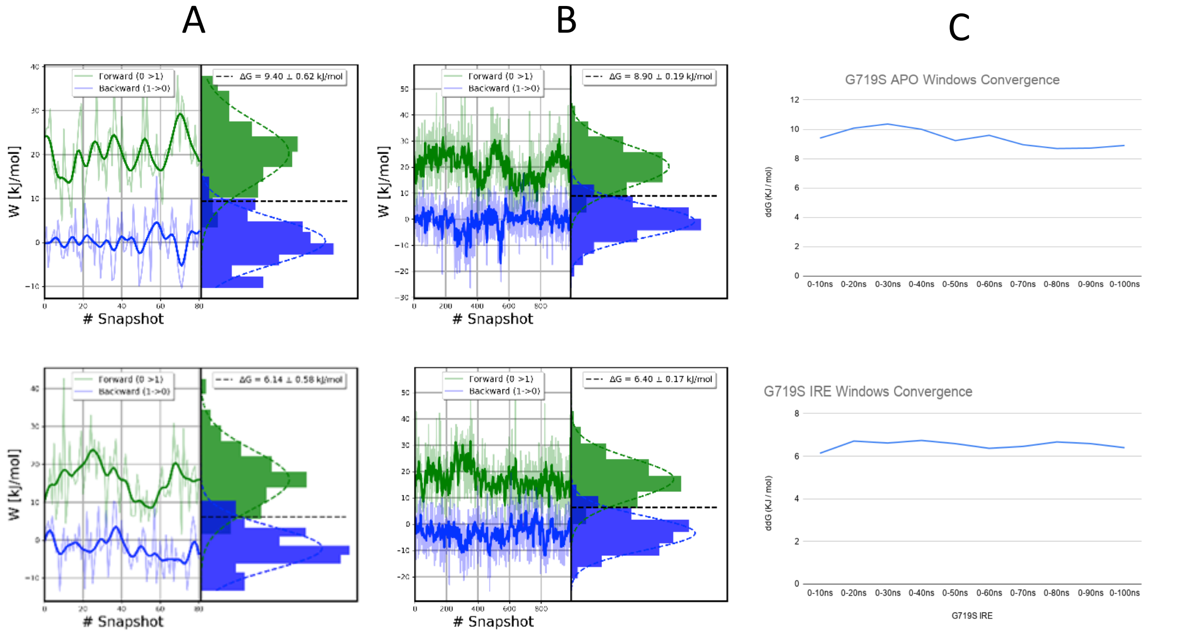


***Suppl. Figure S1:*** *Representative example of convergence in free energy calculations. Top plots correspond to apo and bottom plots to holo EGFR (bound to Gefitinib). Columns A and B correspond to 80x2 and 1000x2 thermodynamic integration runs (forward + reverse) respectively. Column C represents the Bennett Acceptance Ratio (BAR) free energy values obtained with an increasing number of snapshots obtained from the extension of the trajectory.*

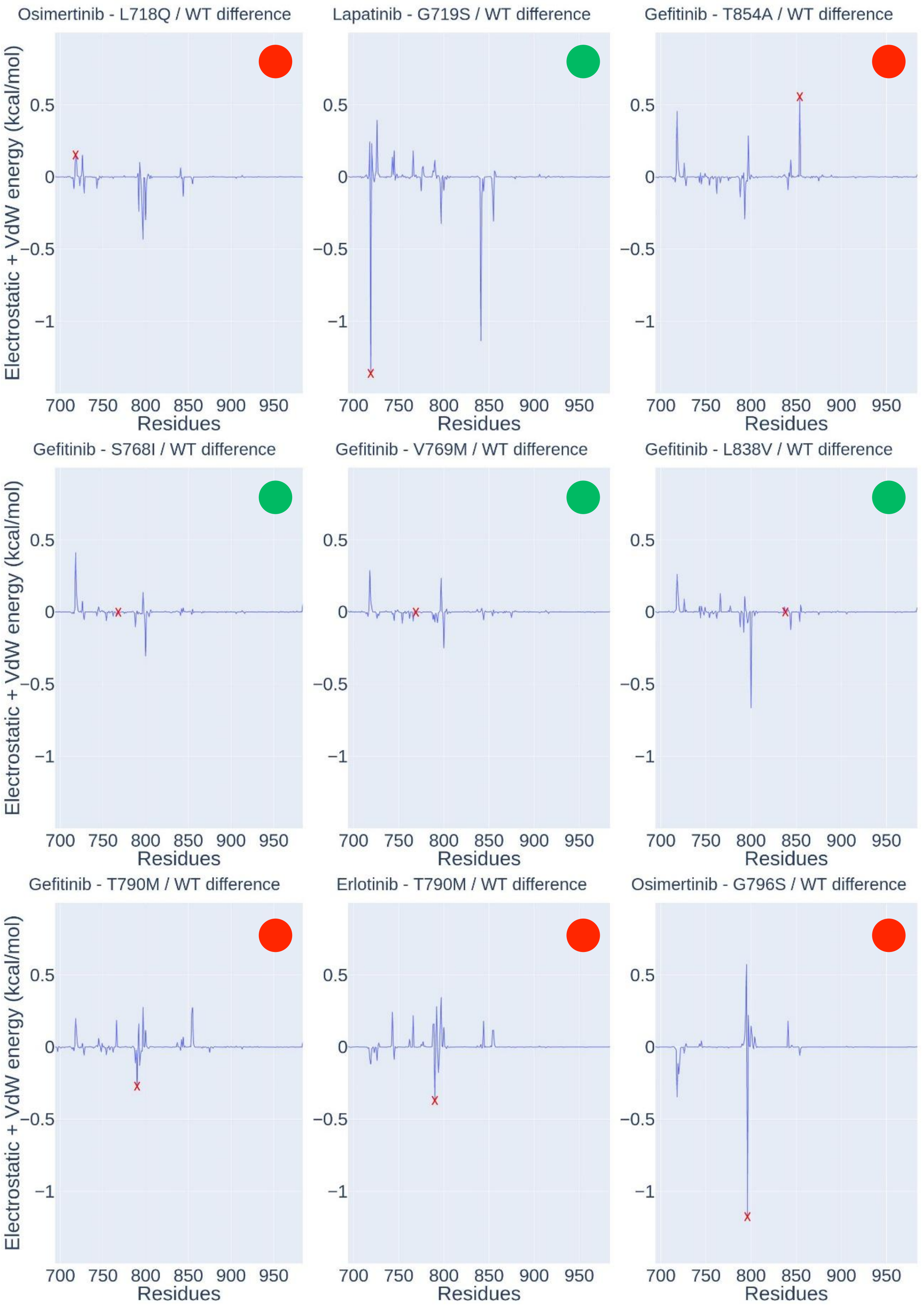


***Suppl. Figure S2****: Example of differential energy maps showing good (first two rows) or bad (last row) predictive power of the impact of mutations on drug binding*. The *red circle means that the mutation is characterized as resistant for the drug considered, and the green circle that the mutated protein remains sensitive to the action of the drug.*

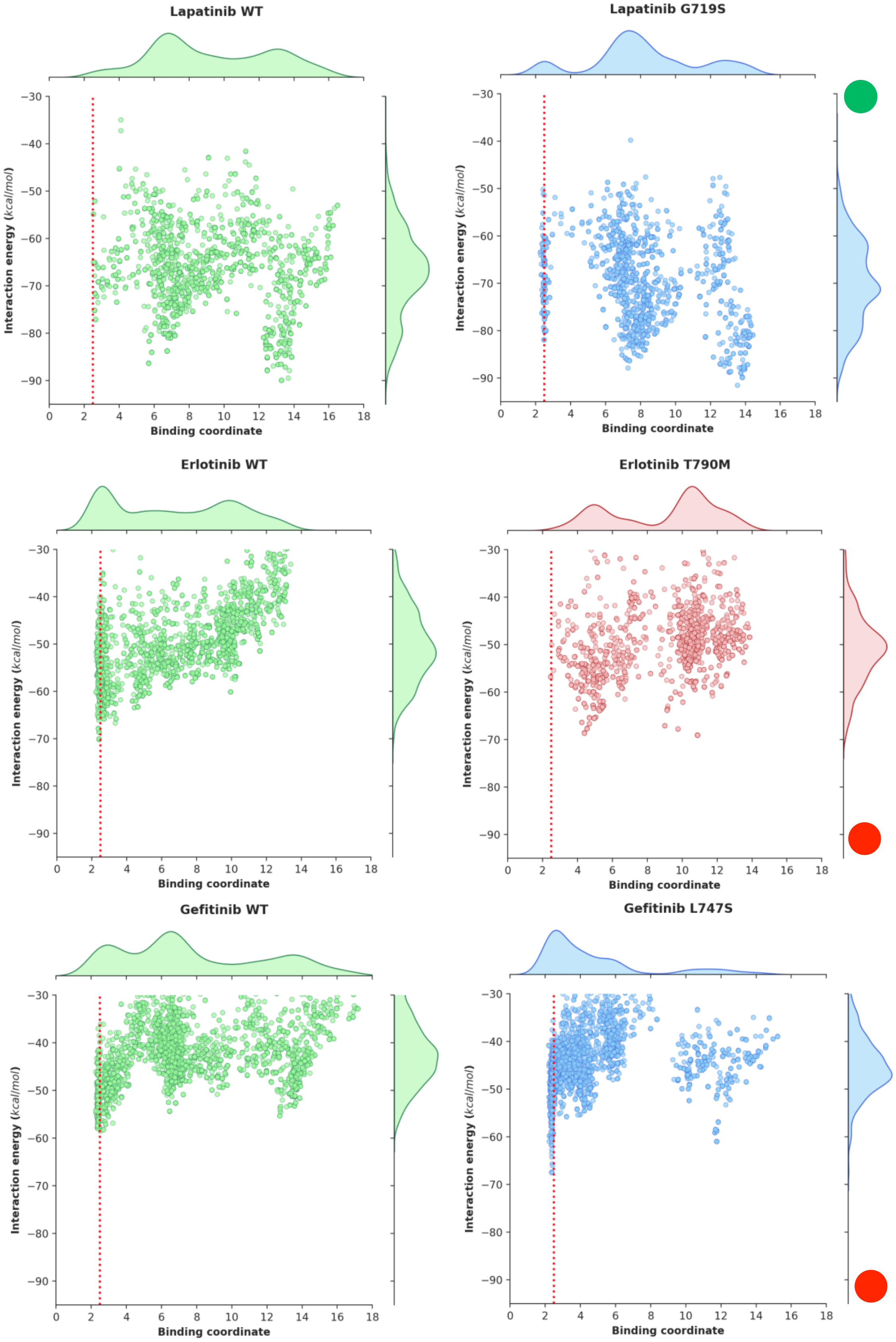
Lapatinib WT
Lapatinib G719S
Erlotinib WT
Erlotinib T790M
Gefitinib WT
Gefitinib L747S
Interaction energy (kcal/mol)
Binding coordinate

***Suppl. Figure S3***. *Examples of successes (top case predicted as sensitive and middle case predicted as resistant) and failures (bottom case erroneously predicted as sensitive) of PELE. Interaction energy at the region of binding determines the affinity of wild type (left) and mutant (right) with a drug. The interaction energy is obtained by subtracting the sum of the energies of the isolated forms of the protein and ligand from the total energy of the system, predicted using the OPLS2005 force field. The binding region is defined as the distance between the backbone carbonyl of Q791 and the hydrogen at carbon-2 of the quinazoline ring of the ligands.*
